# Prehension Kinematics in Humans and Macaques

**DOI:** 10.1101/2021.11.09.467947

**Authors:** Yuke Yan, Anton R. Sobinov, Sliman J. Bensmaia

**Affiliations:** Committee on Computational Neuroscience, University of Chicago, Chicago, IL; Department of Organismal Biology and Anatomy, University of Chicago, Chicago, IL; Neuroscience Institute, University of Chicago, Chicago, IL

## Abstract

Non-human primates, especially rhesus macaques, have been a dominant model to study sensorimotor control of the upper limbs. Indeed, human and macaques have similar hands and homologous neural circuits to mediate manual behavior. However, few studies have systematically and quantitatively compared the manual behaviors of the two species. Such comparison is critical for assessing the validity of using the macaque sensorimotor system as a model of its human counterpart. In this study, we systematically compared the prehensile behaviors of humans and rhesus macaques using an identical experimental setup. We found human and macaque prehension kinematics to be generally similar with a few subtle differences. While the structure of the preshaping postures is similar in humans and macaques, human postures are more object-specific and human joints are less intercorrelated. Conversely, monkeys demonstrate more stereotypical grasping behaviors that are common across all grasp conditions and more variability in their postures across repeated grasps of the same object. Despite these subtle differences in manual behavior between humans and monkeys, our results bolster the use of the macaque model to understand the neural mechanisms of manual dexterity in humans.

**New and newsworthy:** Macaques have been a dominant animal model to study the neural mechanisms of human dexterity because they exhibit complex manual behavior. We show that the kinematics of prehension – a critical dexterous behavior – are largely similar in humans and macaques. However, human preshaping postures are more object-specific and the movement of human digits are less correlated with each other. The thumb, index, and wrist are major driver of these interspecies differences.

## Introduction

The human hand is a remarkably precise and versatile manipulative organ, enabling interactions with objects that range from the mundane – like grasping an object – to the extraordinary – epitomized by virtuosic pianism. Manual dexterity is made possible by the complex anatomy of the hand – which comprises 27 bones articulated by 52 muscle actuators – and by expansive neural circuits involved in motor control and sensing (Sobinov and Bensmaia, 2021).

Our understanding of the neural mechanisms that mediate manual dexterity has relied heavily on experiments with non-human primates, and in particular macaques, whose hands and sensorimotor systems are similar to their human counterparts (Kivell et al., 2016; Sobinov and Bensmaia, 2021; Strick et al., 2021). Indeed, the anatomy of human and macaque hands is largely similar and homologs to the neural structures that support the sensorimotor control of the hand in humans have been identified in macaques.

Qualitatively, manual behavior is also similar in humans and monkeys. Like humans, monkeys pre-shape their hand before grasp in an object-specific way and are capable of some degree of finger individuation (Jeannerod, 1988; Thakur et al., 2008). In both human and monkeys, hand pre-shaping is so object-specific that the identity of an object can be accurately inferred from the (pre-contact) hand posture (Santello and Soechting, 1998; Santello et al., 1998; Schieber and Santello, 2004; Yan et al., 2020). However, to our knowledge, human and monkey manual behavior has never been compared quantitatively using approaches that have been used to characterize manual behavior of either species separately. Careful comparison of human and monkey manual behavior will provide a further assessment of the validity of the macaque sensorimotor system as a model of human manual behavior.

With this in mind, we systematically compared the prehensile behavior of humans and rhesus macaques using an identical experimental paradigm. In brief, humans and monkeys grasped a set of objects handed to them by a robot while we monitored the time-varying joint angles of their hands using a computer vision algorithm. We then compared the hand kinematics of humans and monkeys using a variety of quantitative approaches. We found that, while hand kinematics of prehension are similar in humans and monkeys, these differ in subtle but systematic ways.

## Methods

### Behavioral task

All procedures were approved by the Institutional Review Board of the University of Chicago. Two rhesus macaques were trained to perform a grasping task while seated and with a head fixed in place (Figure 1). Animals were required to keep their arm stationary on armrests between trials, as monitored by a photosensor embedded in the armrest. In the beginning of a trial, an object was attached to a robotic arm (MELFA RV-1A, Mitsubishi Electric, Tokyo, Japan) using a magnet. The arm then presented an object to the monkey with a constant speed always stopping in the same position. While the object was approaching, the animal pre-shaped its hand and grasped it without removing the arm from the photosensor. After 1-3 s, the robotic arm retracted, and the animal had to hold the object with enough force to overpower the magnet. This paradigm ensured that the animal would grasp the object well, instead of just resting a hand on it.

**Figure 1.**
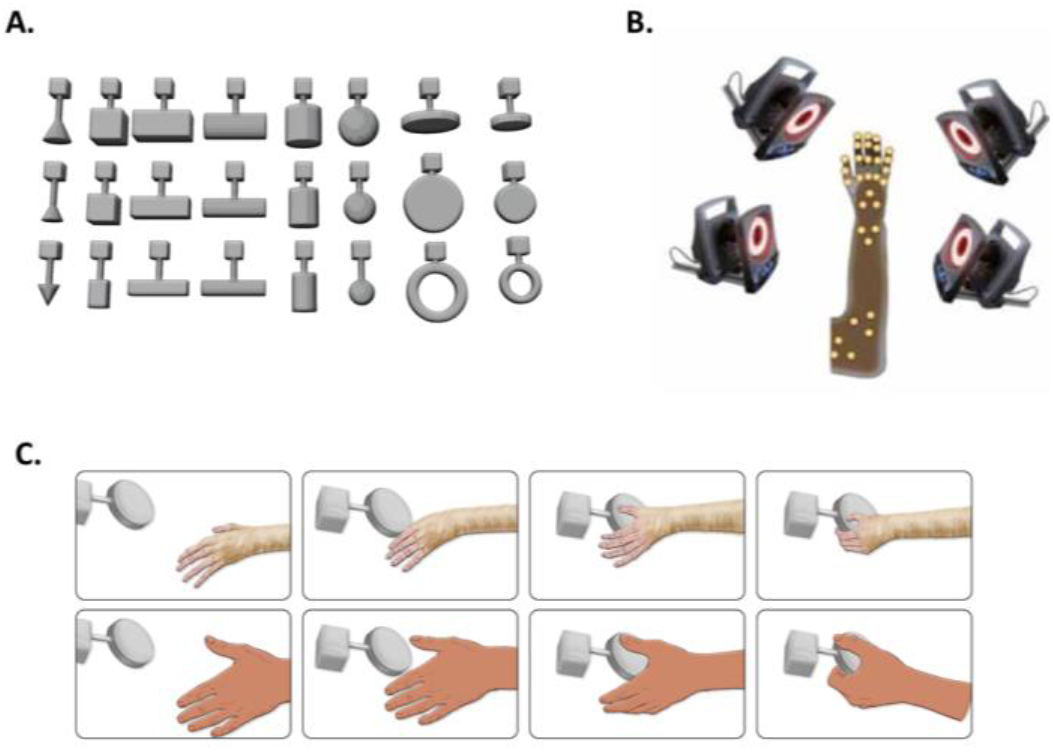
Experimental workflow and grasped objects. A| Overview of apparatus and grasped objects. B| Positions of the tracked markers on macaque forearm. C| Example trial for human and macaque subjects.

Twenty-five different shapes were presented, some of which were used in different orientations, totaling 35 different objects. Each object was presented 8-11 times in a single experimental session. If the animal failed to hold onto the object or lifted its arm, the trial was aborted. For details on the experiment, see (Goodman et al., 2019).

Two right-handed adult human subjects (male and female, 23 and 28 years old) performed the same grasping tasks as the rhesus macaques during a single experimental session. The humans were instructed to grasp the objects tightly without moving their arms off the armrest, but they were not required to hold onto the objects as they retracted.

We compared the kinematics of the two human subjects performing the grasping task to those of two other human subjects performing a reach-to-grasp task with a set of everyday objects (Yan et al., 2020). We found that most analyses yielded similar conclusions with the two data sets (**Supplementary Figure 1, Supplementary Figure 2, Supplementary Figure 3**)

### Recording and processing of kinematic data

We used an infrared motion tracking system to record macaque hand movements (MX T-Series, Vicon, Los Angeles, CA). Thirty-one reflective markers were placed on the bony landmarks of the animal’s arm and hand and tracked in 3D by 14 cameras at a sampling rate of 100 Hz. Each marker was labeled to a corresponding anatomical location using the Vicon Nexus software (Nexus, Vicon, Los Angeles, CA). For details on the data acquisition, see (Goodman et al., 2019).

We used a machine learning method to track the kinematics of the human subjects (Greenspon and Sobinov, 2021). The movements were recorded by 8 video cameras (Blackfly S USB3 BFS-U3-16S2C-CS, FLIR Integrated Imaging Solutions, Inc.) at a spatial resolution of 1440×1080 and a sampling rate of 50 Hz. A machine vision neural network was trained on 80 images per camera per subject to locate 31 manually labelled bony landmarks distributed over arm and hand (Mathis et al., 2018). The network labelled all frames, producing 2D trajectories of the landmarks in the camera plane. We triangulatedthe 2D maker positions using the relative locations of the cameras, to obtain the time-varying 3D positions of the markers.

For both humans and non-human primates, we used OpenSim Inverse Kinematics tool (Seth et al., 2018) to obtain the time-varying joint angles of the hand from the 3D marker trajectories based on a scaled musculoskeletal model of primate hand and forearm (Holzbaur et al., 2005; Saul et al., 2015). We tracked the timing-varying angles of 23 degrees of freedom around 16 joints: flexion-extension, rotation, and radioulnar deviation of the wrist; flexion-extension and abduction-adduction of the carpometacarpal (CMC), metacarpophalangeal (MCP), and interphalangeal (IP) joint of the thumb; flexion-extension and abduction-adduction of the MCP and flexion-extension of the proximal interphalangeal (PIP) and distal interphalangeal (DIP) of each of the four fingers. The angles were smoothed with a 50-ms moving average. We only analyzed kinematics from the start of movement (indicated by the wrist movement from rest) to 100 ms before contact. Joint angles in that period reflect visually guided volitional movements and are not affected by the contact with the objects.

### Subspace similarity

To investigate the similarity in kinematics, we used cross-projection similarity (Todorov and Ghahramani, 2004). In brief, we calculated the amount of variance in the kinematics of one subject that could be explained by the first N principal components (PCs) of a different subject. This value was then normalized to the variance explained by those PCs in the original subject. If the kinematics subspaces were identical for the two subjects, then, the cross-projection similarity would be 1. Note that this metric is not commutative: the projection of kinematics of one subject onto the PCs of the other is not necessarily equal to projection of the kinematics of the second onto the PCs of the first. For that reason, we computed it both ways for each pair of subjects and used their average as the cross-projection similarity.

We first computed the average cross-projection similarity between different human subjects (SH), different monkey subjects (SM), and between pairs of humans and monkeys (SHM) at a fixed dimensionality (9 dimensions, which explains ~90% of the variance). We calculated the average of SH and SM, which represents the mean within-species similarity, and calculated the normalized interspecies difference D = (SH+ SM)/2 – SHM. D is the normalized subspace difference between species. We then calculated the change in D if we remove all joints of a particular digit (e.g., thumb) or a joint group across all digits (e.g., MCP joints). We interpreted a decrease in D upon removal of a digit or joint group as evidence that this digit or joint group contributes to the dissimilarity between monkeys and humans.

### Classification

To quantify the degree to which the hand volitionally conforms to each object, we attempted to classify the grasped objects based on the hand posture 100 ms before contact. To this end, we first calculated PCs from that posture for each subject. Then, we used linear discriminant analysis (LDA) on kinematics projected on all the PCs or on subsets of PCs (Yan et al., 2020), using leave-one-out cross validation: For each object, one trial per object was randomly left out of the training data, and the classifier was tested on those trials. The procedure was repeated with each trial left out. To quantify how much each joint group or digit pre-shapes to the object, we examined classification accuracy using only one digit or joint group.

### Linear dependence of fingers and joint groups

To quantify the degree to which individual joints move independently of other joints, we first regressed the trajectory of each joint on that of every other joint. We then calculated the linear dependence index by averaging all *R*^2^ values of each joint in a digit or joint group (Ingram et al., 2008). Values near 0 denote nearly complete independence and values near 1 denote full linear dependence.

### Variance decomposition

While PCA is informative in identifying axes of maximum variance, it does not provide information about the nature of the variance. To fill in the gap, we employed a modified version of PCA, demixed PCA (dPCA). Briefly, dPCA finds linear dimensions in the data that maximize variance but also identify variance that can be explained by task conditions (Brendel et al., 2011; Kobak et al., 2016). Here, we looked for the variance that could be explained by object identity and trial progression. The latter is a representation of a stereotypical grasping motion, common for all objects, and the rest of the variance was marked as noise. We used the MATLAB package “dpca” by Kobak et al. to perform the dPCA analysis.

## Results

We examined the hand movements (23 joint angles) of two humans and two rhesus macaques as they grasped objects of varying shapes and sizes (**Figure 1**), focusing on the pre-shaping epoch before contact with the object is established. We found, as has been previously shown, that joint kinematics were similar for repeated grasps of the same objects and differed across objects (**Figure 2**). We then set out to quantitatively compare the prehensile behavior of humans and monkeys.

**Figure 2.**
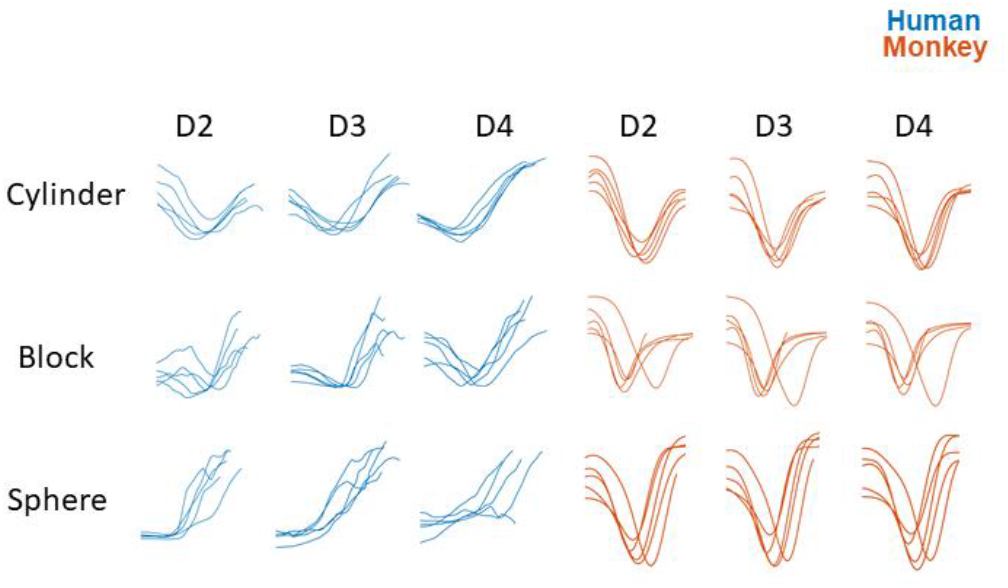
Example kinematics traces. Time varying joint angle traces of Digits 2 through 4 in humans and monkeys. Each row corresponds to a different object and each curve to a different trial. The joint angles shown are MCP flexions of the fingers.

### Basic features of prehension kinematics

First, we examined the parameters of movement for monkeys and humans, including the range of motion, speed, and acceleration of each joint as well as the duration of grasping movements (**Figure 3**). We found that the ranges of motion were similar for monkey and human joints, but the speeds and accelerations differed in joint-specific ways. For example, the distal interphalangeal joints (DIP) tended to accelerate and move faster in humans than monkeys (**Figure 3**) (Mann-Whitney U test, speed: n=11620, p<1e-06; acceleration: n=11620, p<1e-06). The reverse was true for the proximal finger joints (MCP flexion/extension), which tended to accelerate more rapidly and reach higher peak speeds in monkeys. (p<1e-06). Overall, monkeys performed the task more rapidly than did humans (Mann-Whitney U test, n=2324, p<1e-06). Viewed through this lens, humans tend to move their distal joints more than do monkeys, which prefer to complete the movement using proximal hand joints. This difference may be attributed to the more common use of the hook grip in monkeys than humans, which involves the distal joints remaining in a hook configuration during the hand closing.

**Figure 3.**
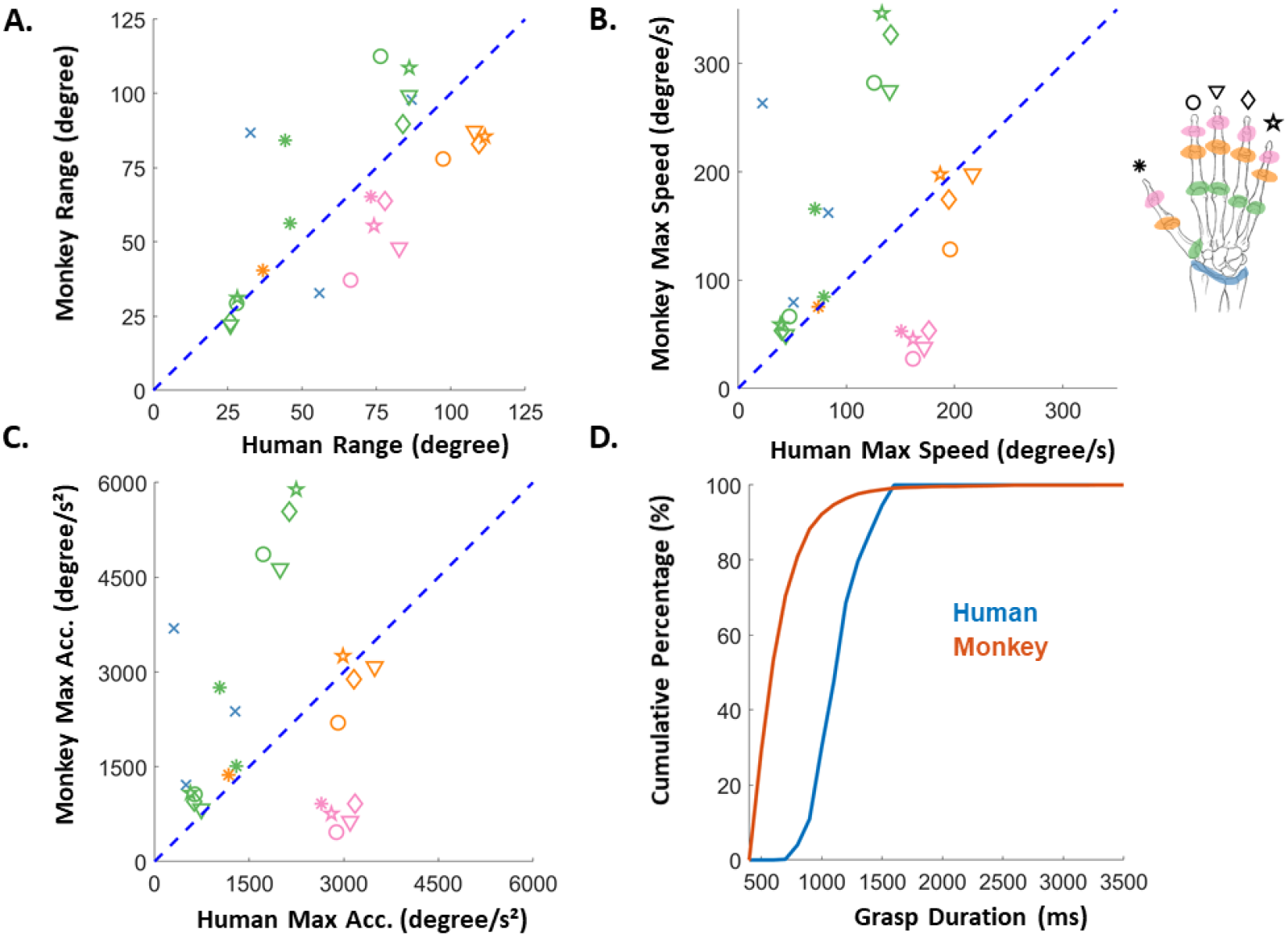
Basic kinematic statistics of grasping motions for monkeys and humans. A| Average range of motion of each joint for monkeys and humans. Joint group is coded by color and digit by marker shape. Note that MCP joints have more data points than others because MCP joints are capable of abduction, whereas DIP and PIP joints are not. B| Scatter plot of the maximum speed of each joint. Speed is averaged across trials and subjects. C| Scatter plot of the maximum acceleration of each joint. D| Cumulative distribution plot of grasp durations of monkeys (red) and humans (blue). Y axis shows the percentage of grasp durations falling below the threshold.

### The structure of prehension kinematics

Next, we examined and compared the structure of manual behavior in the two species. To this end, we performed a principal component analysis (PCA) on the prehension kinematics of monkeys and humans.

We found that human kinematics tended to be more complex as evidenced by a shallower cumulative variance plot: more principal components (PCs) were required to account for the same proportion of variance in human than monkey kinematics (**Figure 4A**). However, this separation was relatively modest, limited to one or two PCs. Furthermore, the first two PCs were remarkably similar for humans and monkeys (accounting for 67% and 77% of the variance, respectively): the first involved changes in hand aperture, the second mainly involved movements of the wrist (**Figure 4B**).

**Figure 4.**
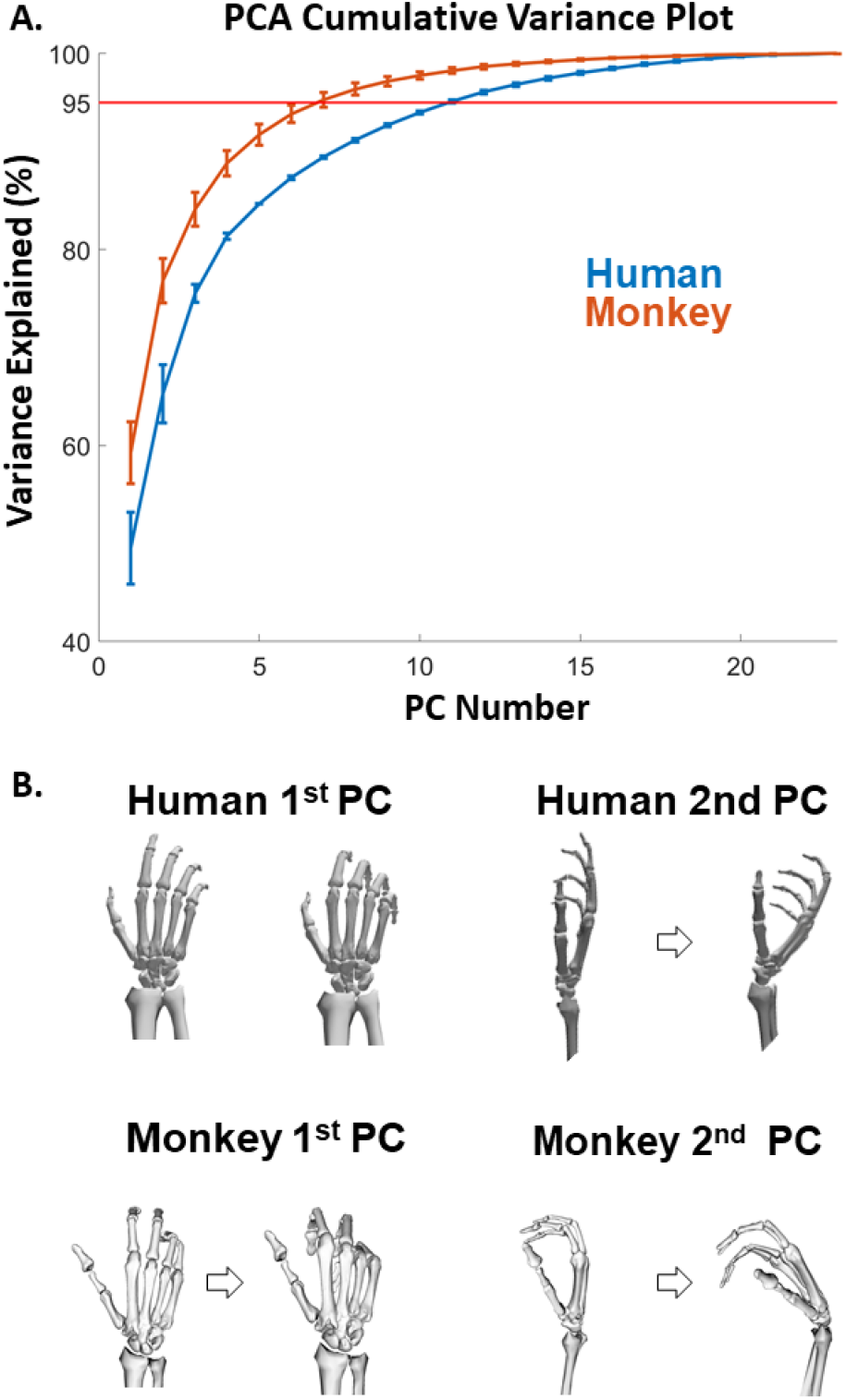
Principal component analysis of kinematics. **A**. Cumulative variance explained as a function of the number of PCs. PCs are arranged in descending order of eigenvalues. **B**. The hand movements produced by the 1^st^ and 2^nd^ PCs in monkeys and humans.

### Dependence of joints

The kinematic subspaces reflect the linear dependence of different hand joints. Indeed, each PC denotes a specific pattern of correlated joint angles. To compare patterns of inter-joint correlations across humans and monkeys, we analyzed the linear dependence of each digit and joint group on the other joints, measured as the mean fit (*R^2^*) of a regression of the time-varying angle of one joint on that of all the other joints. We found that monkeys yielded consistently higher linear dependence across all digits than did humans (**Figure 5A**) (Wilcoxon signed rank test, n = 40, p = 2.53e-04). In other words, monkey digits were significantly less individuated during grasp than were their human counterparts, consistent with the results from the PCA. In both monkeys and humans, the thumb was the most individuated digit, yielding a lower linear dependence index than the other digits (**Figure 5A**). The wrist, MCP, and PIP joints of monkeys were more linearly dependent on the other joints than were their human counterparts (Wilcoxon signed rank test, n = 36, p = 5.36e-04) whereas the individuation of the DIP was similar in the two species (**Figure 5B**) (Wilcoxon signed rank test, n = 10, p=0.813).

Note, further, that the interdependence of joints at the level of joint groups is somewhat idiosyncratic whereas the interdependence of joints grouped by digits is rather consistent (**Supplementary Figure 4**).

**Figure 5.**
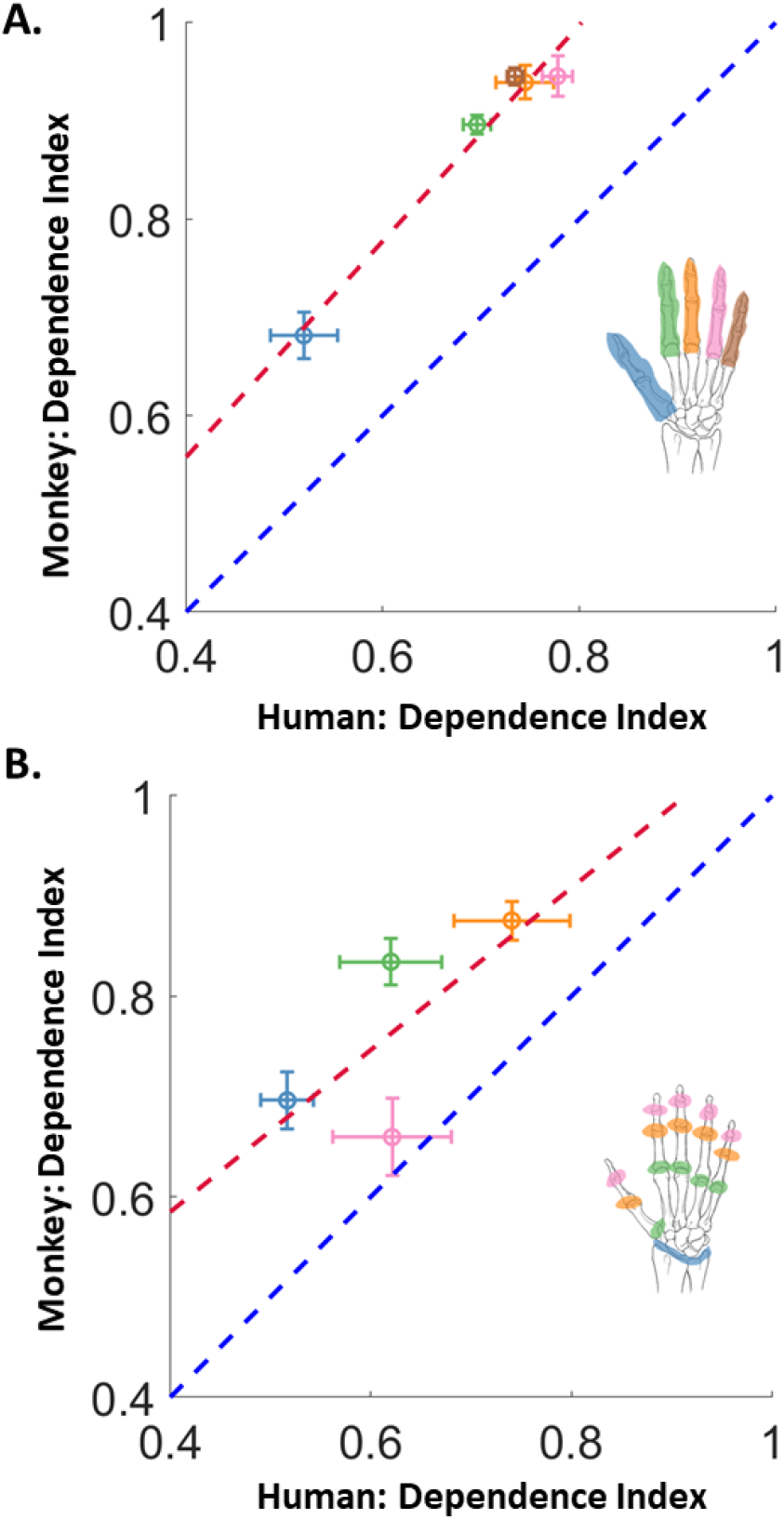
Dependence Index of digits and joint groups. A| Linear dependence of each digit for monkeys vs. humans. B| Linear dependence index of each joint group for monkeys vs. humans. Blue dashed line is the identity line; red dashed line is the regression line.

### Kinematic subspaces of monkeys and humans

Next, we aimed to quantify the similarity of the kinematic subspaces in humans and monkeys. To this end, we calculated how much variance in the hand kinematics of one subject could be accounted for using the PCs of another, human or monkey, a measure known as cross-projection similarity. We found that, on average, the first 10 PCs of one species could account for about 80% of kinematics of the same species and 75% of the variance of the other species (**Figure 6A**), suggesting a high degree of similarity in the structure of human and monkey prehensile kinematics. Nonetheless, the cross-species similarity was significantly lower than its within-species counterpart (human-human vs human-monkey: Wilcoxon signed-rank test, n = 46, p = 3.79e-06, monkey-monkey vs human-monkey: Wilcoxon signed-rank test, n = 46, p = 1.2e-06) (**Figure 6A**).

**Figure 6.**
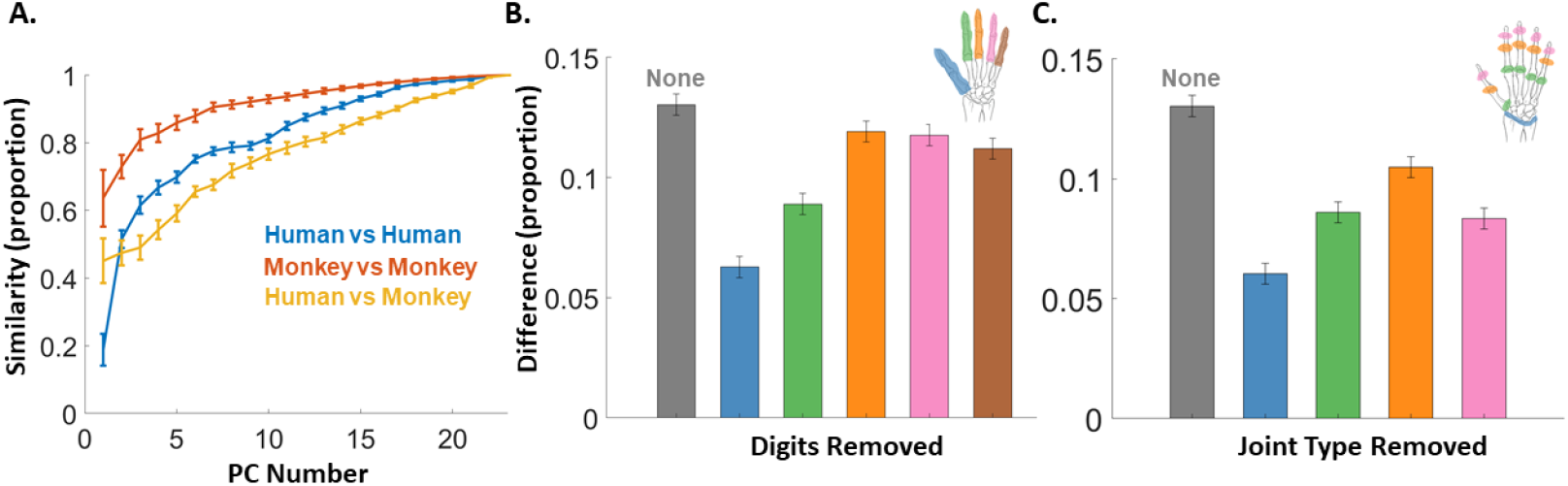
Subspace similarity between humans and monkeys. **A|** Cross-projection similarity (measured in proportion variance explained) as a function of the PCs added. Error bars denote SEM. **B|** Inter-species difference with individual digits removed. The difference is measured as the distance between the human-monkey similarity to the average of the human-human similarity and the monkey-monkey similarity. The bar marked as “None” represents the difference value calculated without removal of any digits. **C|** Inter-species difference with different joint groups removed.

Given the relatively subtle difference in human and monkey kinematics, we set out to examine which digits or joint groups contributed most to this difference. To this end, we computed an “interspecies difference” measure by subtracting the mean across-species similarity from the mean within-species similarity. We then applied this metric after removing a digit (**Figure 6B**) or a joint group (wrist, MCP, PIP and DIP) (**Figure 6C**). To the extent that removing a digit/type led to a decrease in interspecies difference, this digit/joint group contributes to the observed difference between humans and monkeys. We found that all digits/joint groups contributed to the human-monkey difference to a certain degree, with thumb, index, and wrist contributing most (**Figure 6BC**). We repeated these analyses with a different subspace similarity measure – principal angle – and obtained similar results (**Supplementary Figure 5**).

### Complexity of prehensile behavior

One manifestation of dexterity is that manual behavior is tailored to the task. The object specificity of pre-contact hand postures during prehension is an example of such task specificity. With this in mind, we compared the object dependence of human and monkey pre-shaping postures and examined how this dependence was distributed over the joints. To these ends, we used linear discriminant analysis to classify the grasped object based on the hand posture 100 ms before contact, when the hand pre-shaped to the object but before volitionally adopted hand posture is distorted by object contact. We found that, for both monkeys and humans, objects could be classified with high accuracy based on hand posture. Both human subjects exhibited hand pre-shaping postures that were highly object-specific (**Figure 7A**). However, the degree of object specificity was very different for the two monkeys: One yielded classification performance commensurate with that of humans, the other exhibited pre-shaping hand kinematics that were far less object-specific, yielding lower classification performance.

**Figure 7.**
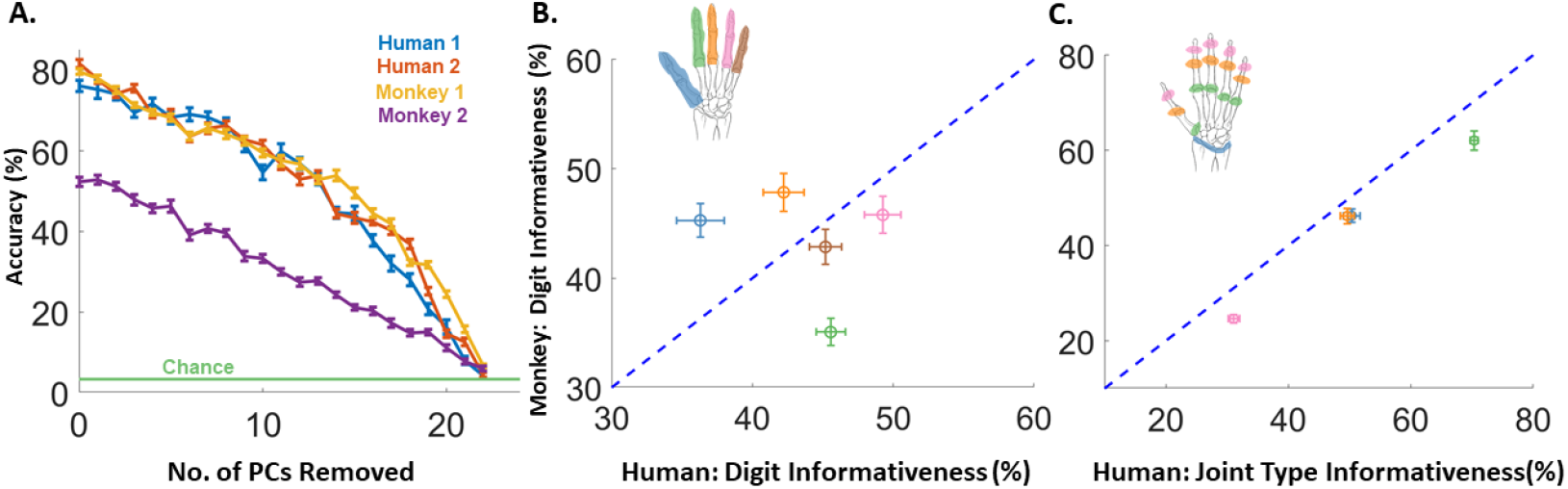
Object specificity of hand preshaping. **A|** Object classification performance based on postures measured 100 ms before object contact, with an increasing number of high-variance PCs removed. **B|** Classification accuracy based on the kinematics of each digit individually for monkeys vs. humans. **C|** Classification accuracy based on the kinematics of each joint group individually for monkeys vs humans.

We then assessed how this object-specificity was distributed over the space of hand postures in two ways. First, we measured classification performance as we gradually removed high-variance PCs. Preservation of classification after removal of PCs suggests that prehensile behavior occupies a high-dimensional space (Yan et al., 2020). In both humans and monkeys, we found that object information was distributed over a large number of PCs (**Figure 7A**)(cf. (Yan et al., 2020)). Even after removal of all but 1 PC, classification performance was above chance (~4% > 2.9%). Second, we examined the degree to which different joint groups adopted object-specific postures. To this end, we calculated the accuracy of object classification based on the postures of subset of joints, grouped joints by digit (**Figure 7B**) or type (**Figure 7C**). Again, humans and monkeys yielded similar results, with subtle differences. For example, the index finger was less informative in monkeys than humans (Mann-Whitney U test, n=50, p=3.35e-02). Contrary to expectation, the thumb was much more informative in monkeys than in humans (Mann-Whitney U test, n=50, p=3.54e-02). The object specificity of the joint groups, however, was consistent across humans and monkeys, with the MCP being the most informative and DIP the least (**Figure 7C**). Note, however, that these indices of informativeness also depend on the objects, as evidenced by the fact that they differ in human subjects across different objects sets (**Supplementary Figure 3**). Thus, prehensile behavior in both humans and monkeys occupies a high-dimensional kinematics space and is distributed over joint groups in similar ways.

As mentioned above, the hand postures of one monkey were far more object-specific than were those of the other. In contrast, the two humans exhibited highly similar prehensile behaviors (**Figure 7A**). In light of this, we repeated the above analyses on the kinematics of individual subjects to assess the degree to which our conclusions were skewed by the observed differences across monkeys. We found that basic prehensile features and linear dependence indices were consistent across monkeys (**Supplementary Figure 6**, **Supplementary Figure 7**, **Supplementary Figure 4**), despite these significant differences in the object-specificity of hand pre-shaping (**Supplementary Figure 8**, **Supplementary Figure 9**).

### Decomposition of hand kinematics variance

Finally, we sought to understand the determinants of human and monkey prehensile kinematics. To this end, we used demixed PCA (dPCA) (Brendel et al., 2011; Kobak et al., 2016) to decompose the kinematics of each subject into components that were stereotypical across objects (time-dependent), components that were specific to individual objects (object-dependent), and components that varied across repeated presentations of the same object (noise). We found that monkey kinematics comprised more time-dependent and noise variance and less object-dependent variance than did their human counterparts (**Figure 8**). In other words, monkey prehensile postures were less object-specific, more stereotyped, and noisier than were their human counterparts. Furthermore, the decreased object-specificity observed in the classification analysis for Monkey 2 (**Figure 7A**) is driven by greater noise in this monkey’s prehension kinematics than in those of the other monkey or in humans (**Supplementary Figure 9**).

**Figure 8.**
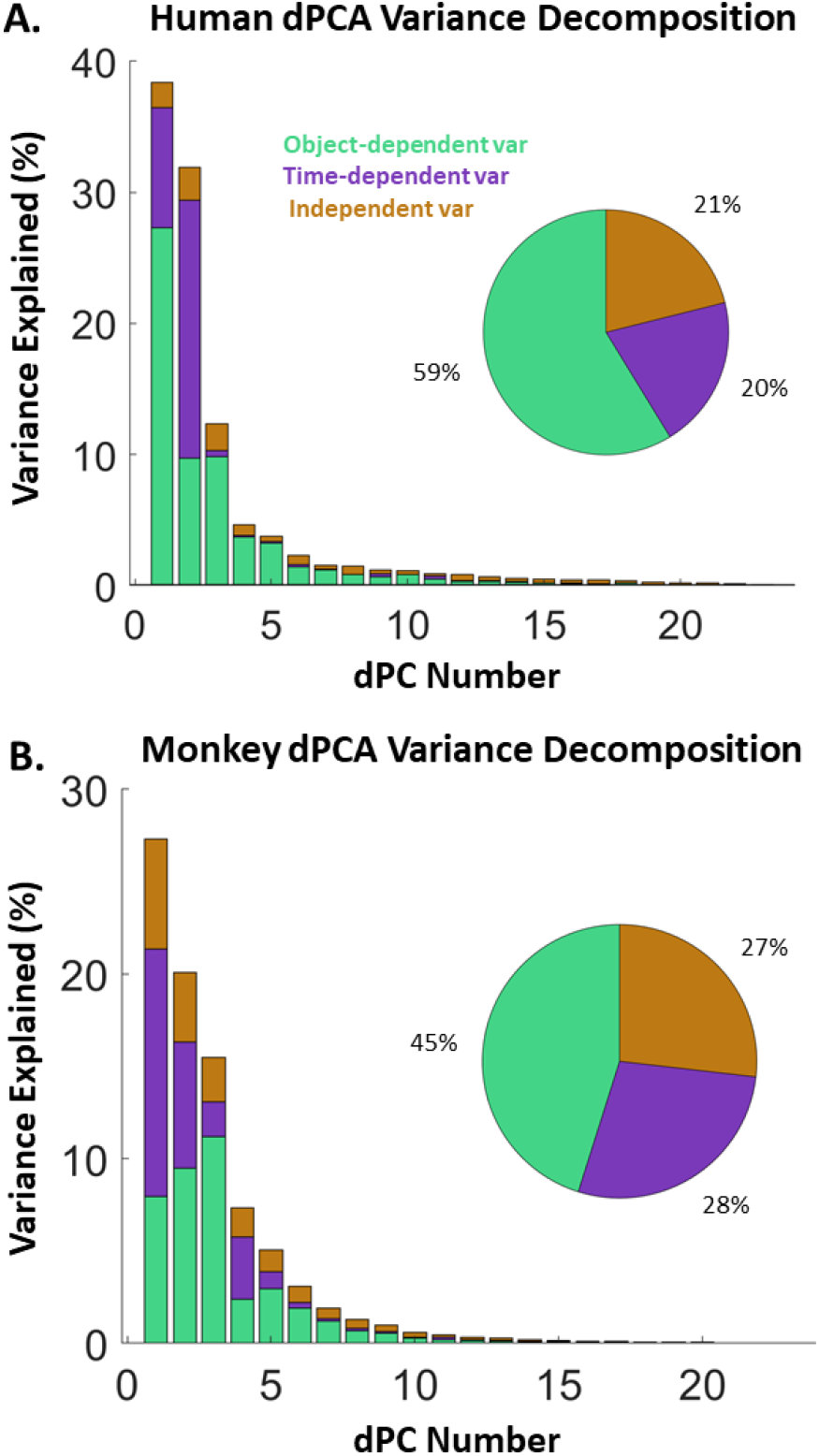
dPCA analysis and variance decomposition. A. Human variance decomposition. Bar plots shows the variance explained by each dPC arranged in descending order of eigenvalues. For each bar, the contribution by each type of variance is shown by color. Pie chart shows the overall proportion of each type of variance. B. Monkey variance decomposition.

## Discussion

In summary, we find that, while the basic statistics of prehensile kinematics – effective ranges of motion, speeds, and accelerations – differ somewhat between humans and monkeys, the structure of manual behavior is similar for the two species. Both humans and monkeys exhibit high-dimensional pre-shaping kinematics, though monkey kinematics are more stereotyped and noisier than are their human counterparts.

### Kinematic similarities and differences

Among non-human primates, macaques arguably possess the most human hands except for the (non-human) apes (Kivell et al., 2016). Macaque hands are endowed with a semi-opposable thumb, share much of the musculature with humans, and can individually move their fingers (Schieber, 1991; Häger-Ross and Schieber, 2000; Diogo and Wood, 2012). Human and macaque hands differ most strikingly in the respective morphologies of their thumbs. First, the macaque thumb is much shorter than is the human one in comparison to the index and middle fingers (Napier, 1967; Patel and Maiolino, 2016), putatively to prevent it from interfering with arboreal locomotion. Second, the pronounced saddle shape of the trapezium bone supporting thumb carpometacarpal joint in humans produces considerable rotation of the thumb as it abducts, presenting its distal pad to that of the fingers, allowing greater opposability (Kivell, 2016). Third, more muscles articulate the human than the macaque thumb (Diogo and Wood, 2012). All of these differences contribute to increased manipulative ability of humans, exemplified by tool-making and strong precision grip (Young, 2003; Kivell et al., 2011; Rolian et al., 2011; Karakostis et al., 2021; Williams-Hatala et al., 2021). We found that the thumb was the most independent digit in both humans and monkeys and the thumb was a significant contributor to the dissimilarity between human and monkey prehensile kinematics.

The index finger also significantly contributed to the difference in kinematics, possibly due to its higher individuation in humans, enabled by the anatomy of the *extensor indicis* muscle. In humans, this muscle only extends the index finger whereas, in macaques, it also extends the middle finger (Diogo and Wood, 2012). Interestingly, the kinematics of the index finger are the least object-specific in macaques whereas the kinematics of the thumb are the least object-specific in in the humans. The greater stereotypy in the thumb and index may reflect the fact that these digits contribute the most to grasp. As macaques commonly use hook grasps, the index finger provides stability; as humans favor the power grip, the thumb plays the equivalent stabilizing role.

Grasping is commonly characterized by measuring hand aperture, commonly defined as the distance between the tips of index and thumb (Castiello and Dadda, 2019). Macaques change their aperture with higher velocity and acceleration than humans during reach and grasp tasks (Christel and Billard, 2002). We found that this difference in aperture dynamics is produced by the difference in the movement of the proximal finger joints (Figure 3ABC), an effective strategy as proximal joint deflections result in larger endpoint deflections.

Often overlooked, the wrist is the most complex joint of the hand, connecting up to 17 bones in primates and transferring the load between hand and arm for locomotion and manipulation (Kivell, 2016). Functionally, the carpal bones of non-human primates can lock together for support during terrestrial locomotion or allow for a high range of motion during other behaviors. In humans, however, the wrist is not used for locomotion, which has led to the fusion of some wrist bones, a decrease in bone volume, and changes to its microanatomy, making the human wrist somewhat simpler anatomically, although better suited for manipulation and tool making (Williams et al., 2010; Marzke, 2013; Schilling et al., 2014; Bird et al., 2021). One might thus expect large differences in wrist kinematics. Indeed, we found human wrist movements to be more independent than their macaque counterparts, and differences in wrist kinematics were a major contributor to interspecies differences in prehensile behavior.

### Control of volitional movements

In primates, large swaths of cortex are dedicated to hand control, much larger than in other species (Penfield and Rasmussen, 1950; Baldwin et al., 2018; Lemon, 2019; Mayer et al., 2019; Kaas et al., 2020; Strick et al., 2021). The motor cortex of many primates – mostly Old World monkeys and apes (including humans) – sends projections directly onto spinal motoneurons and thus has more access to the muscles than does the motor cortex of other mammals (Bernhard et al., 1953; Heffner and Masterton, 1975; Iwaniuk et al., 1999; Rathelot and Strick, 2009). That prehensile kinematics occupy a high-dimensional space in both humans and macaques likely reflects these expansive and specialized neural circuits that mediate manual dexterity in humans and macaques (Sobinov and Bensmaia, 2021). Indeed, the representation of hand posture in primary motor cortex is higher dimensional than the already high dimensional hand postures themselves (Schaffelhofer et al., 2015; Okorokova et al., 2019; Yan et al., 2020).

Macaques and humans also possess highly developed cortical fields in posterior parietal cortex (PPC) that convert visual information about the object into a motor plan (Mountcastle et al., 1975; Tunik et al., 2005; Schaffelhofer and Scherberger, 2016; Kaas et al., 2020). Specifically, the anterior interparietal area is crucial for the visuomotor transformation from shape to grip type (Taira et al., 1990; Sakata et al., 1995; Durand et al., 2007; Gharbawie et al., 2011; Stepniewska et al., 2014). These mechanisms of visuomotor transformation are likely the substrate that underlies the ability of humans and macaques to pre-shape their hands to objects to be grasped and, more generally, to exhibit manual behaviors that are appropriate for specific interactions. In both humans and macaques, a large proportion of the variance in hand movements was object-specific and, though this proportion was higher in humans than monkeys, differences were dwarfed by similarities. Rats and mice have a comparatively small PPC and are incapable of these sophisticated visuomotor transformations (O’Connor et al., 2021).

## Conclusions

We show that prehensile behaviors in humans and monkeys are very similar when examined through a detailed, quantitative lens. This similarity likely reflects the similarity in end effectors and associated neural circuits. We expect that such detailed behavioral comparisons across multiple species may help disentangle links between neural mechanisms and behavior.

## Acknowledgements

This work was supported by NINDS grant NS122333.

## Supplementary Figures

**Supplementary Figure 1.**
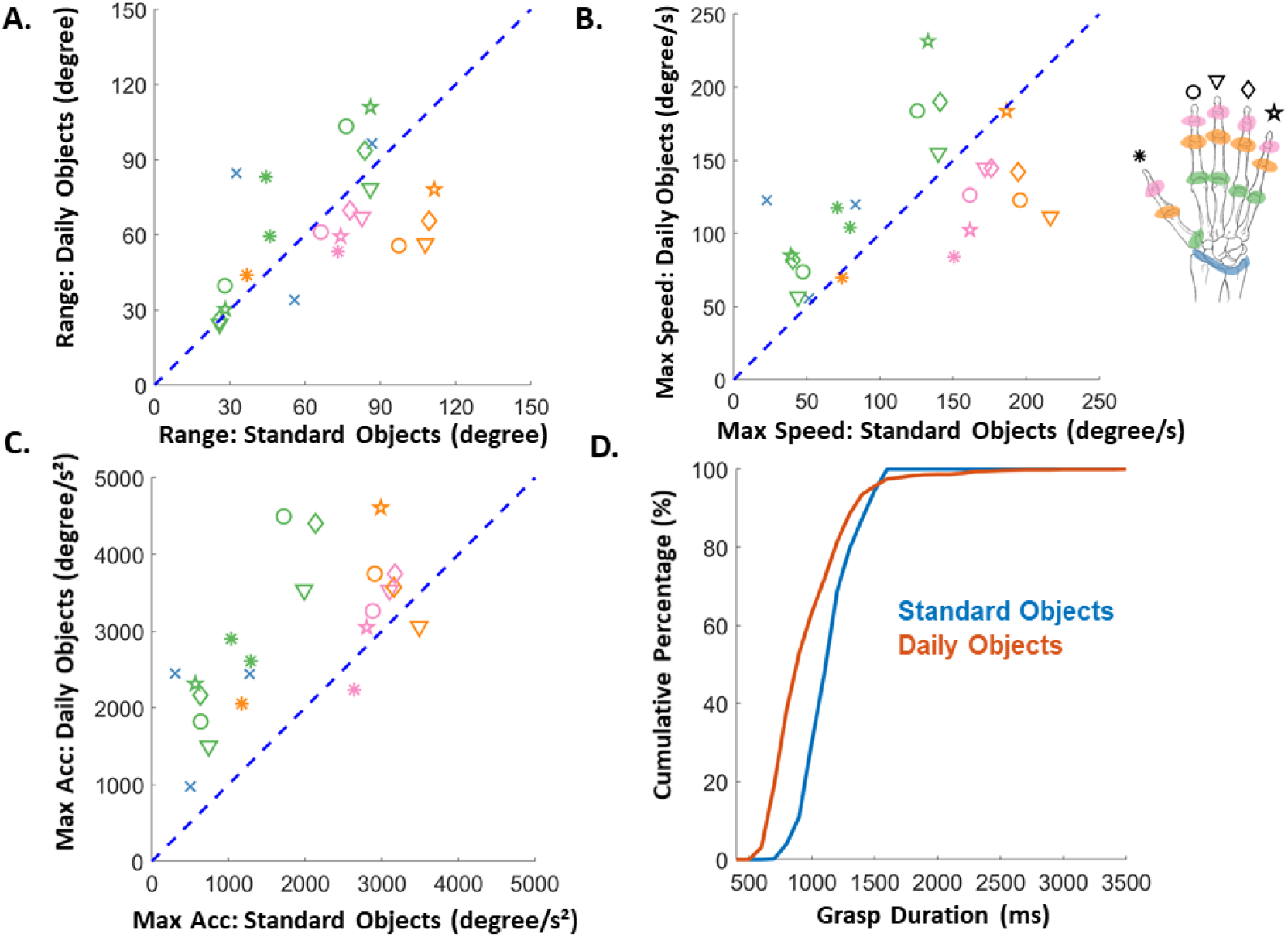
Basic Kinematics Statistics: Two different object sets. A| Average range of motion of each joint for daily objects and standard objects (the same objects used in the monkey experiments). Different colors denote different joint groups and different markers denote different digits (see the top right diagram). B| Scatter plot of the maximum speed of each joint. Speed is averaged across trials and subjects. C| Scatter plot of the maximum acceleration of each joint. D| Cumulative distribution plot of grasp durations of monkeys (red) and humans (blue). Y axis shows the percentage of grasp durations falling below the threshold. The daily objects were grasped using a standard reach to grasp paradigm whereas the standard objects were handed to the subject with a robot. Despite these differences in objects and paradigms, hand kinematics were similar in the two conditions.

**Supplementary Figure 2.**
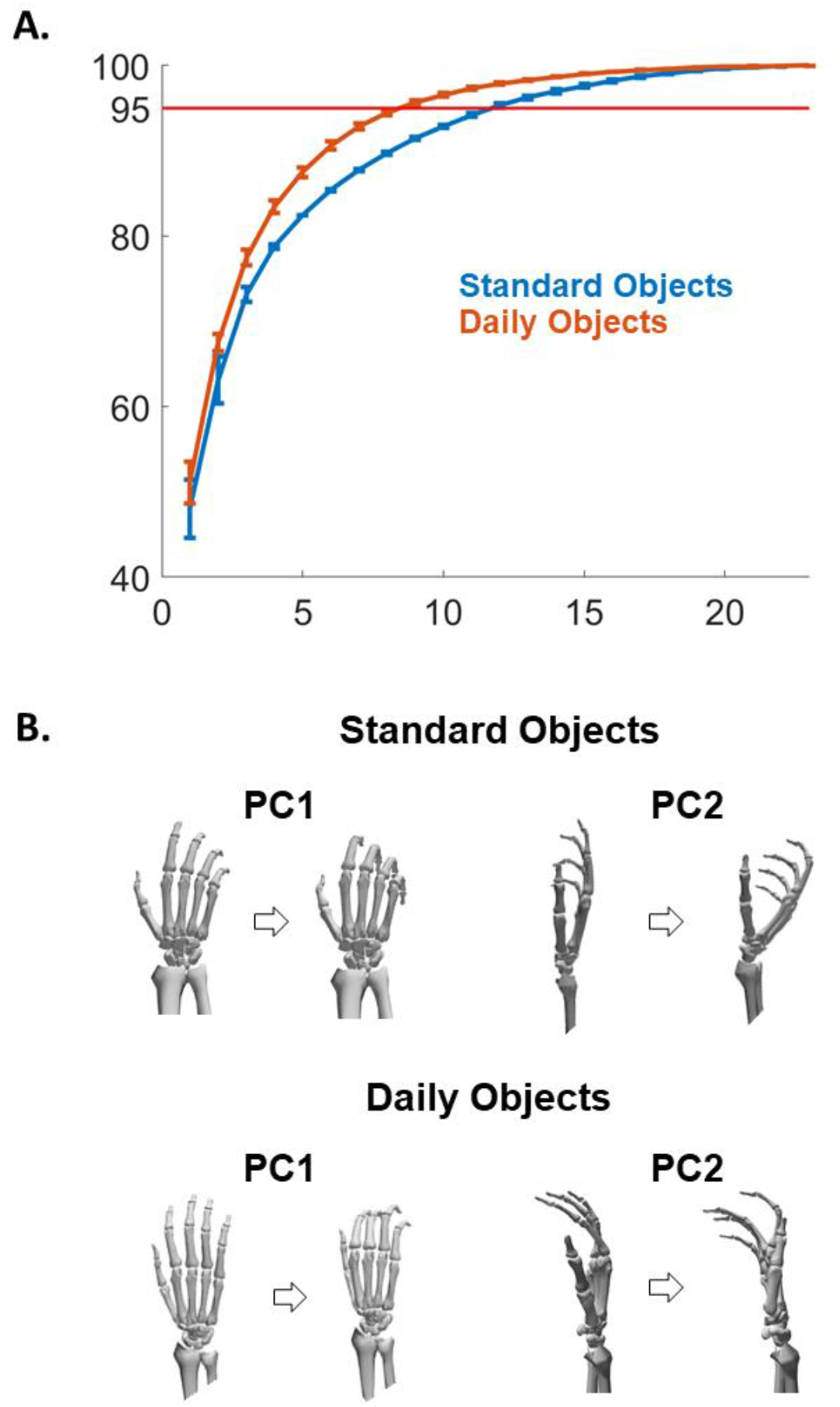
PCA Analysis of Kinematics: Two object sets. **A**| Cumulative variance explained as a function of the number of PCs. PCs are arranged in descending order of eigenvalues. Object set is color coded. **B|** The hand movements produced by the 1^st^ and 2^nd^ PCs of the same human subject for two sets of objects. The daily objects were grasped using a standard reach to grasp paradigm whereas the standard objects were handed to the subject with a robot. Despite these differences in objects and paradigms, hand kinematics were similar in the two conditions.

**Supplementary Figure 3.**
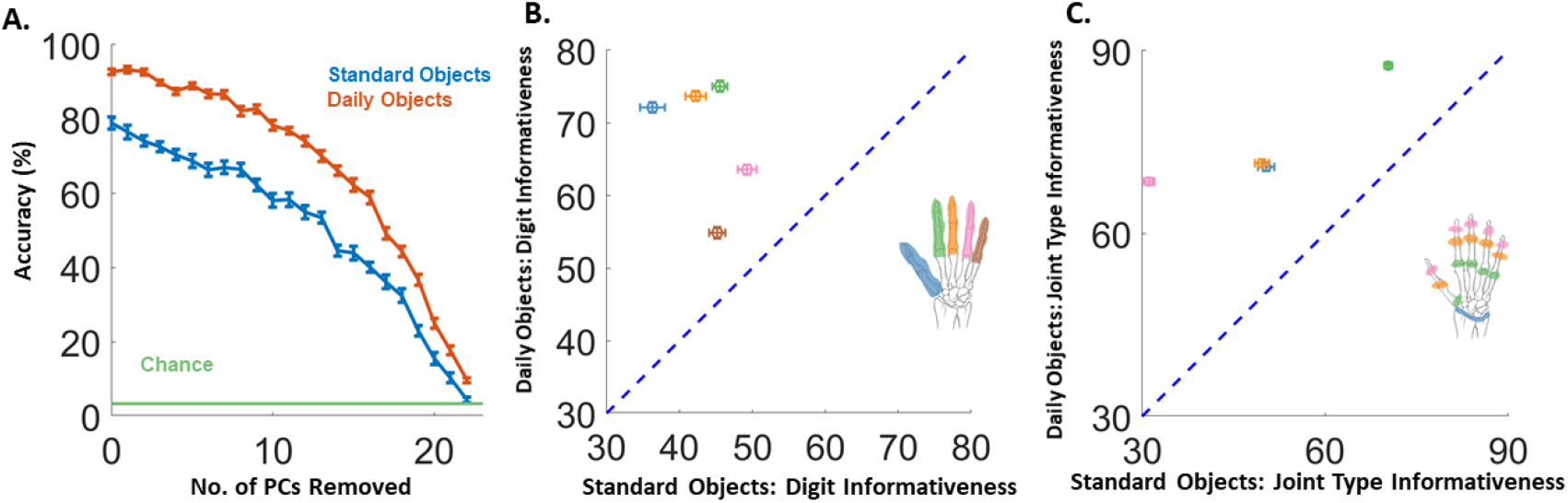
Classification accuracy and joint informativeness across two object sets. **A**| Mean accuracy of object classification for humans with increasing number of high-variance PCs removed. **B**| Classification performance based on the kinematics of each digit separately. **C**| Classification performance based on the kinematics of each joint group separately. Classification of the daily objects was better than that of standard objects, likely because of the greater diversity of the former.

**Supplementary Figure 4.**
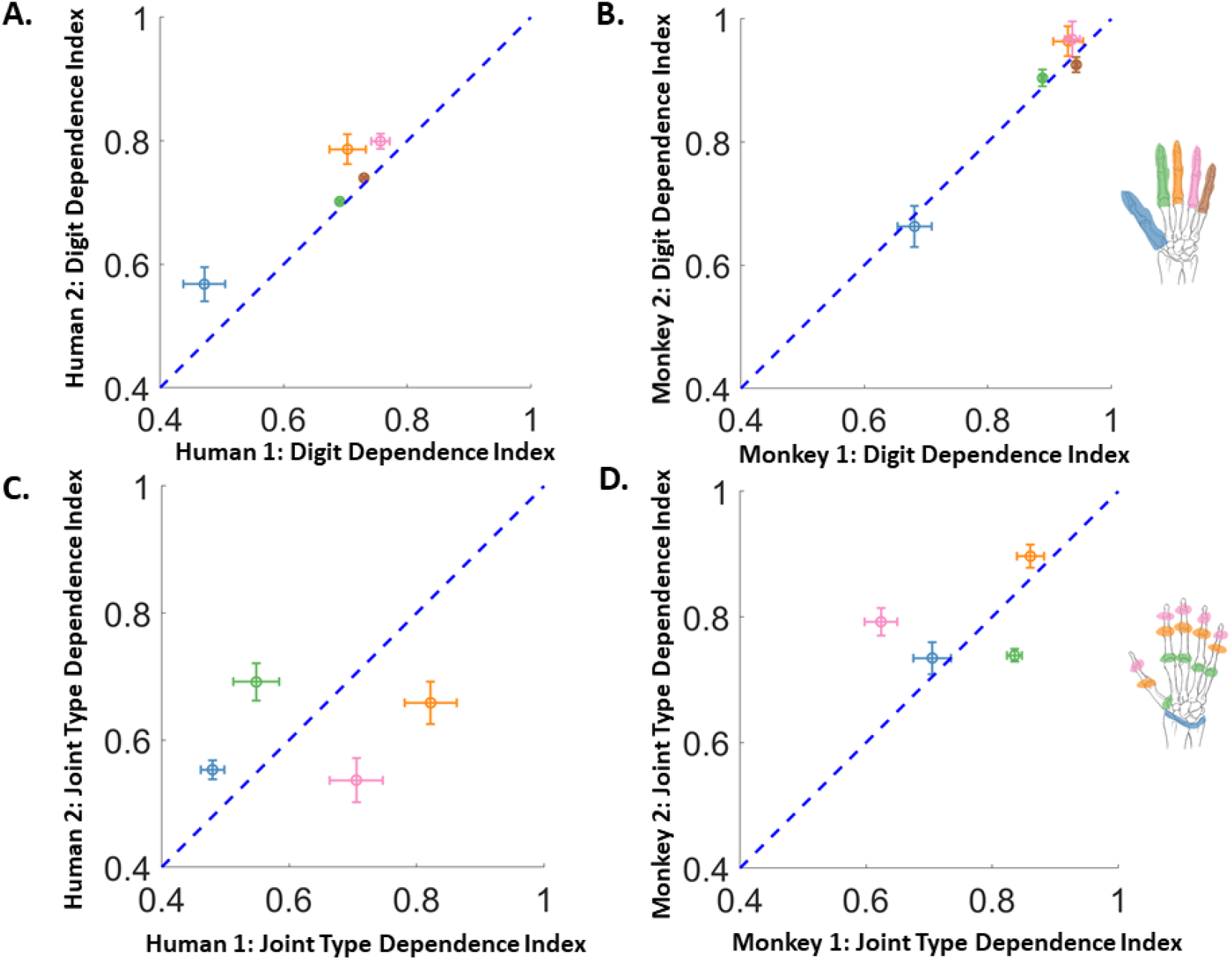
Joint Linear Dependence: Individual Subjects. **A|** Scatter plot of the linear dependence of each digit. **B|** Same as panel A but for Monkey Subject 1 vs Monkey Subject 2. **C**| Scatter plot of the linear dependence of each joint group for Human 1 vs Human 2. **D|** Same as panel C but for Monkey 1 vs Monkey 2.

**Supplementary Figure 5.**
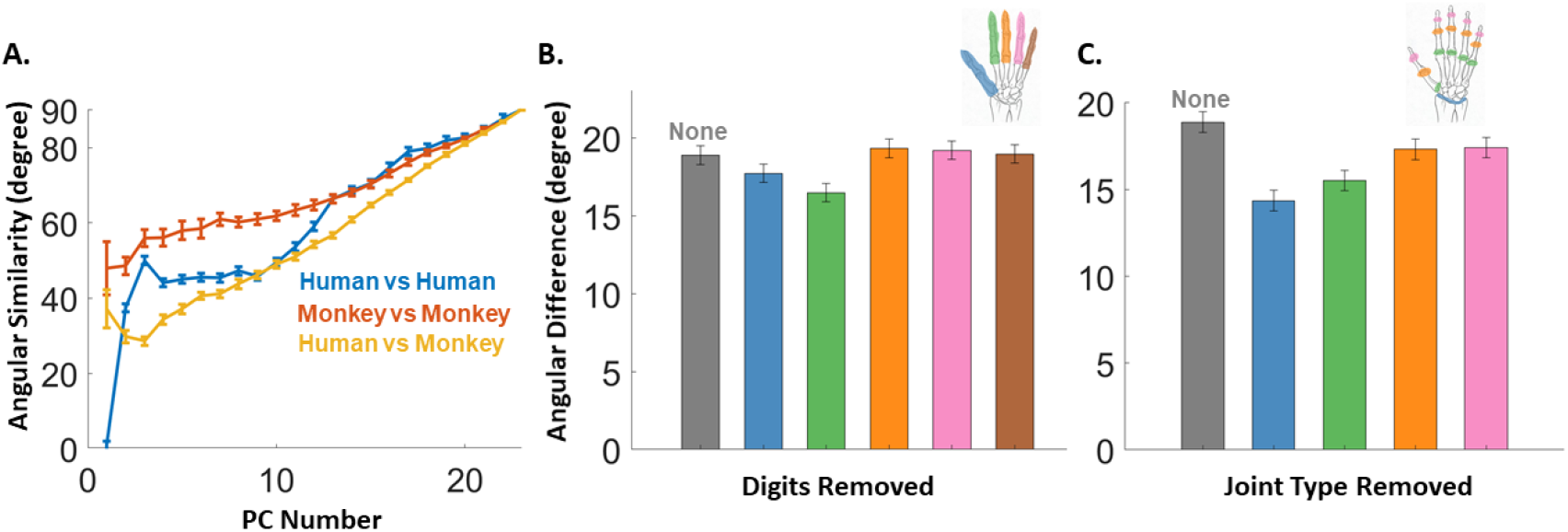
Subspace similarity measured by principal angle. We repeated the analysis in **Figure 6** with a different subspace similarity measure: the principal angle. A| Angular similarity (90 – avg. principal angle, higher means more similar) between species or within species. B| Angular difference as we remove different digits from the data. Digits are color coded and grey bar represents the subspace difference when no digit was removed. C| Angular difference as we removed different joint group from the data. The results are similar to those obtained with cross-projection similarity, except that the thumb and index are reversed in their respective contributions to the dissimilarity.

**Supplementary Figure 6.**
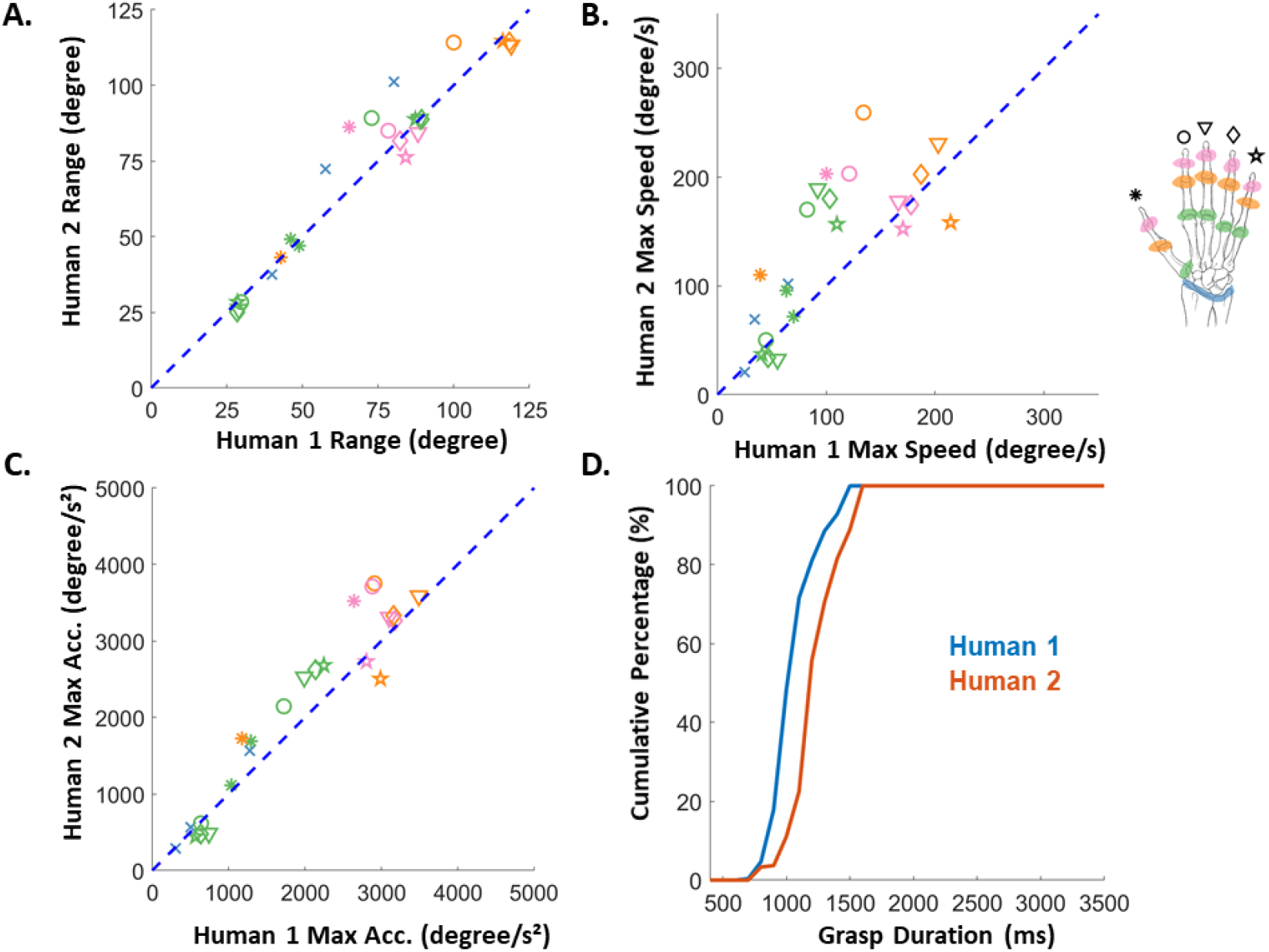
Basic Kinematics Statistics: Human 1 vs Human 2. A| Average range of motion of each joint for human 1 vs human 2. B| Scatter plot of the maximum speed of each joint. Speed is averaged across trials and subjects. C| Scatter plot of the maximum acceleration of each joint. D| Cumulative distribution plot of grasp durations of monkeys (red) and humans (blue). The two human subjects exhibited very similar kinematics.

**Supplementary Figure 7.**
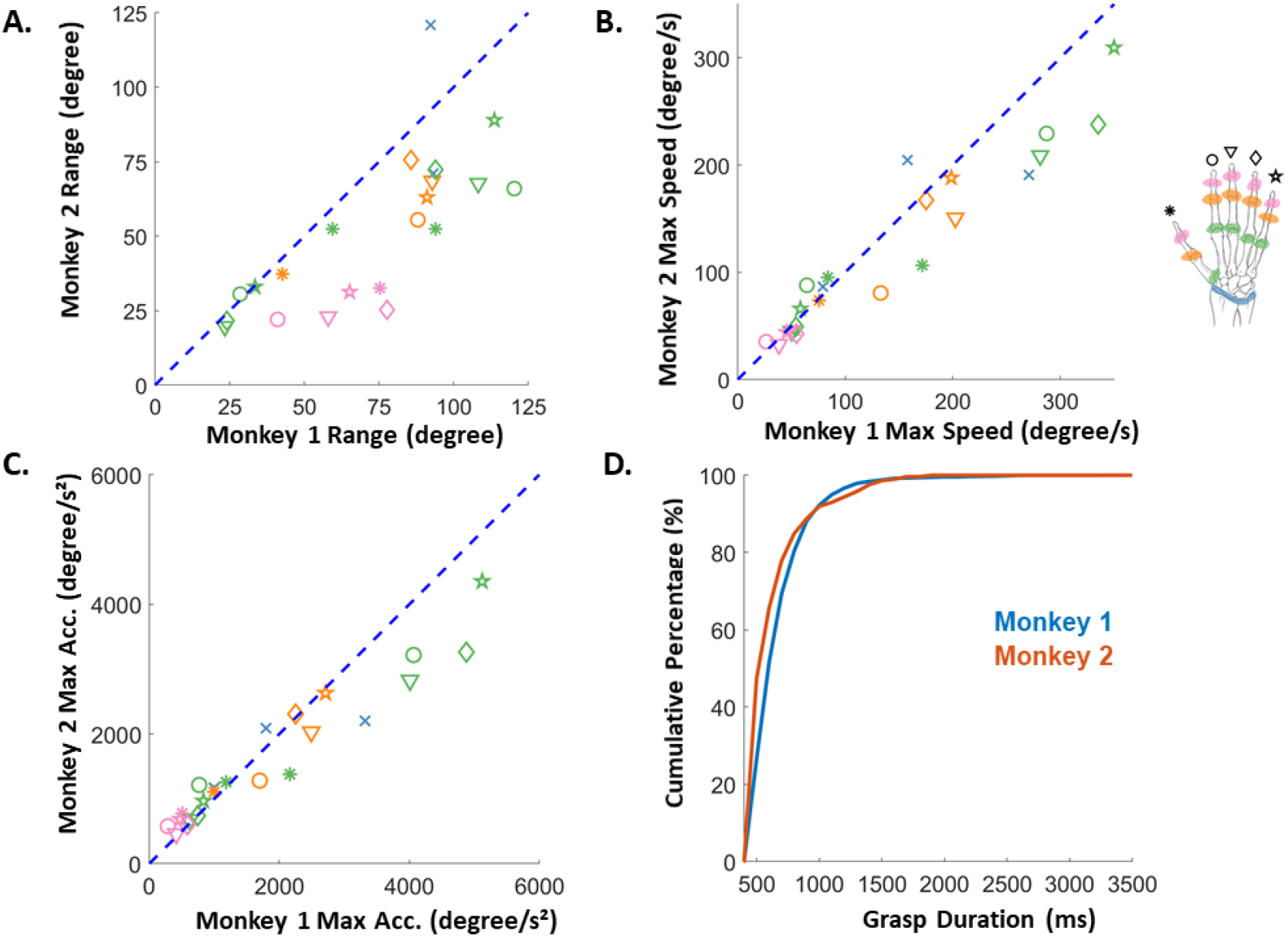
Basic Kinematics Statistics: Monkey 1 vs Monkey 2. A| Average range of motion of each joint for monkey subject 1 vs monkey subject 2. B| Scatter plot of the maximum speed of each joint. Speed is averaged across trials and subjects. C| Scatter plot of the maximum acceleration of each joint. D| Cumulative distribution plot of grasp durations of monkeys (red) and humans (blue). The two monkey subjects exhibited very similar kinematics.

**Supplementary Figure 8.**
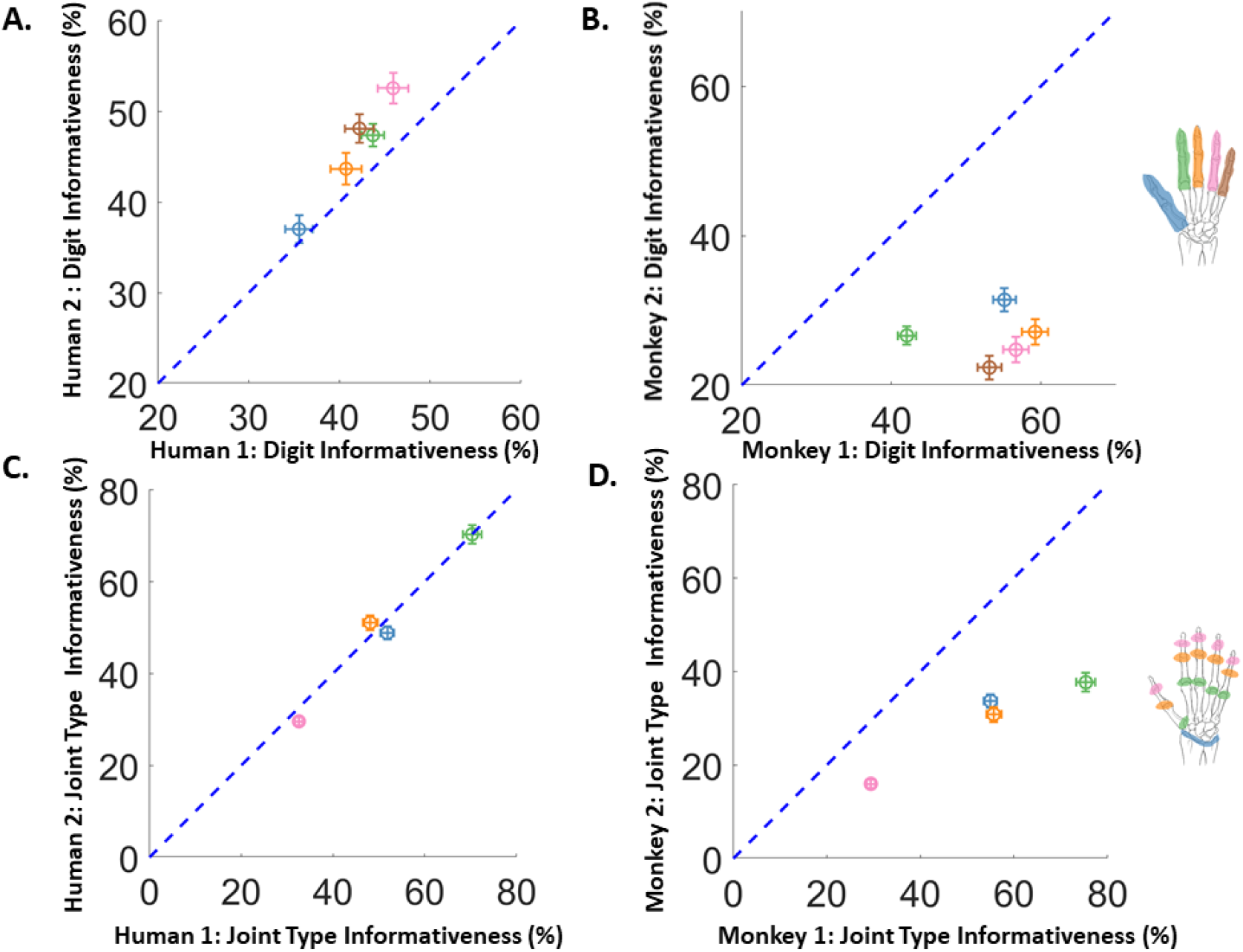
Joint Informativeness: Individual Subjects. **A|** Scatter plot of the informativeness of each digit in classification, measured by the classification accuracy of grasped objects if using that digit alone. **B|** Same as panel A but for Monkey Subject 1 vs Monkey Subject 2. **C|** Scatter plot of the informativeness of each joint group for Human 1 vs Human 2. Joint group is color coded as in the hand diagram. **D|** Same as C but for Monkey 1 vs Monkey 2.

**Supplementary Figure 9.**
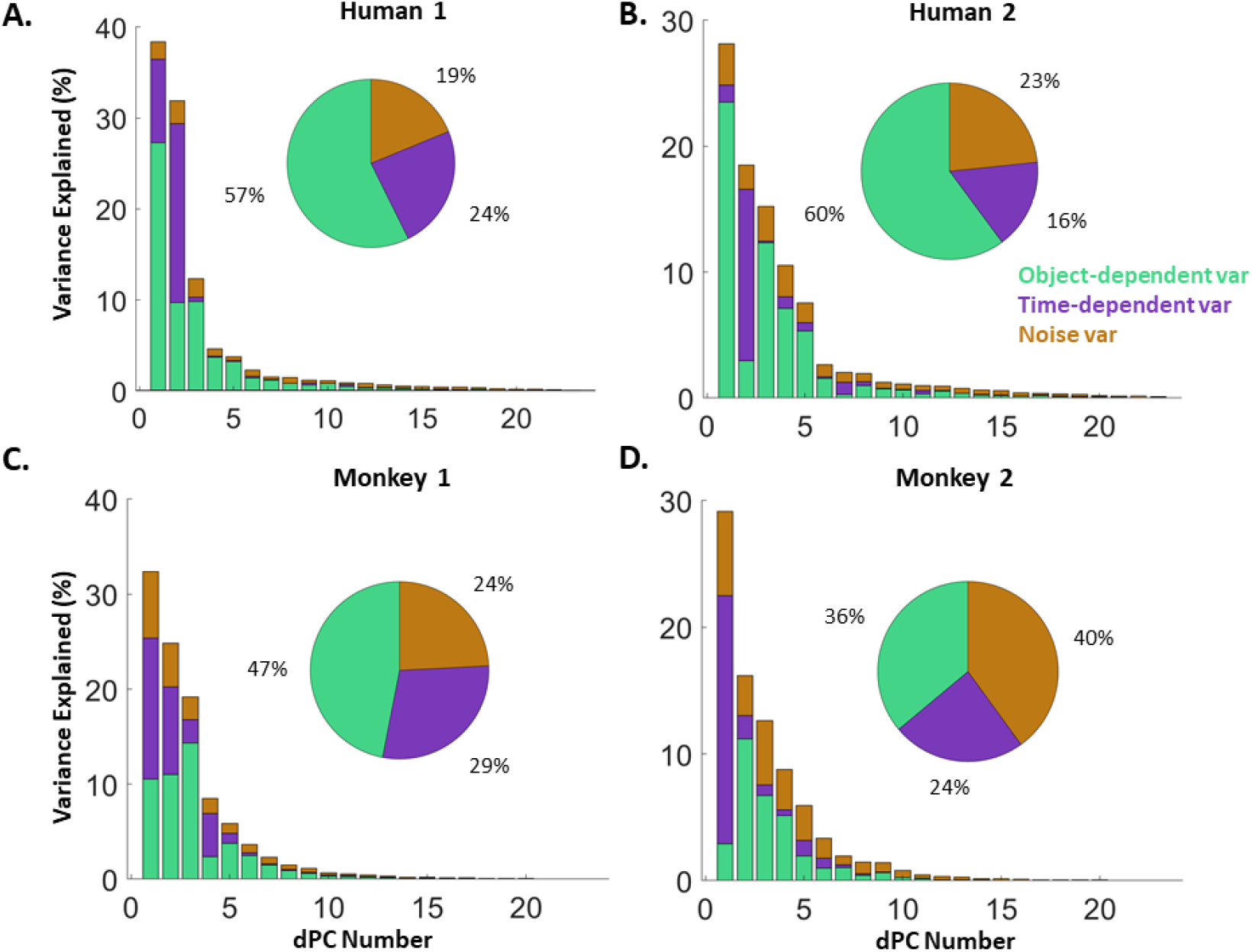
dPCA analysis and variance decomposition: Individual Subjects. **A|** Variance decomposition for human 1. **B|** Variance decomposition for Human 2. **C|** Monkey 1. **D|** Monkey 2

**Supplementary Figure 10.**
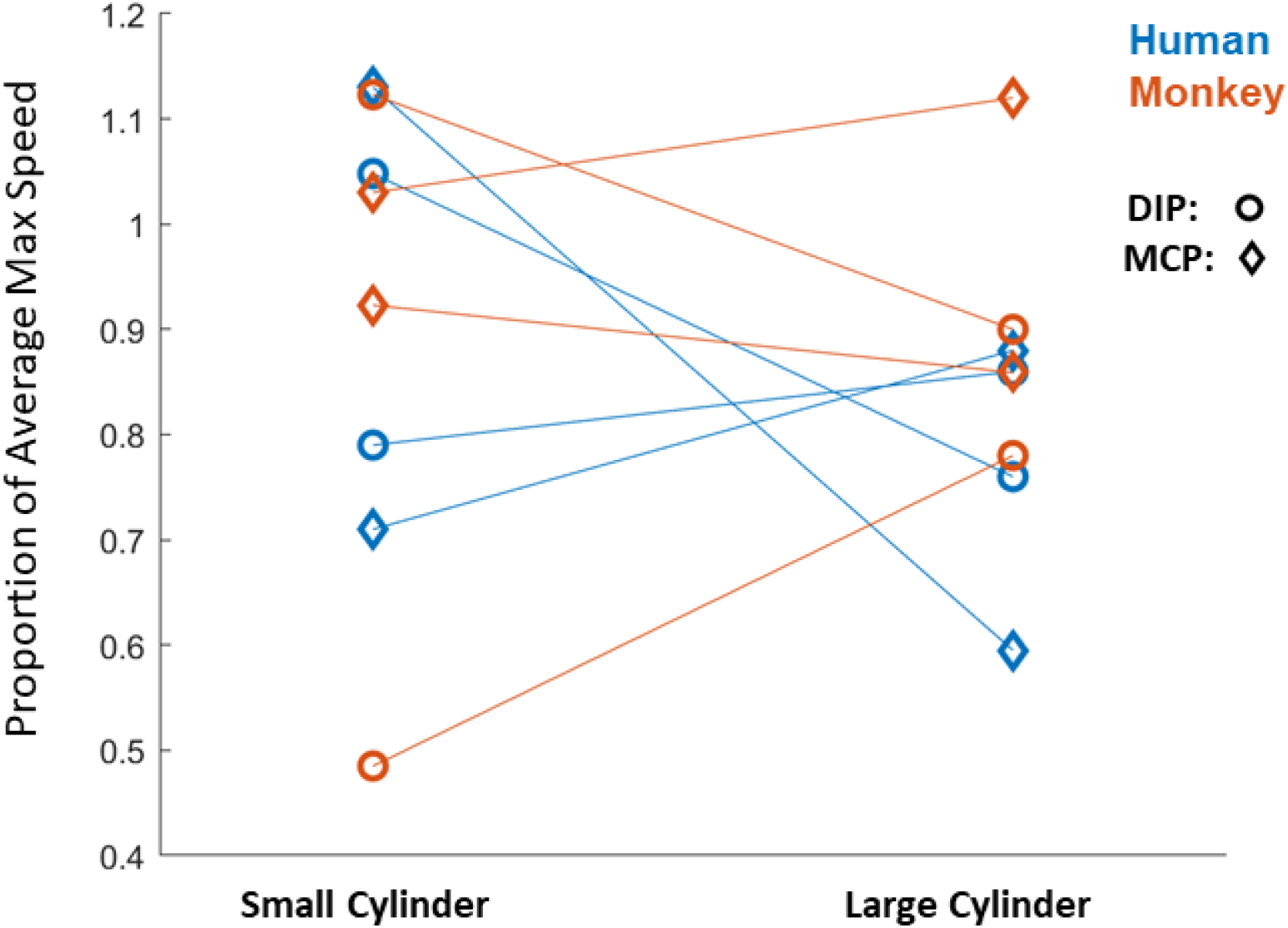
Joint Max Speed vs Object Size. Scatter plot of the joint max speed (as a proportion of the average max speed across all objects) for the small vs. large cylinder. Species are color coded and joint groups are coded by marker symbol, as shown on the top right. Repetition of the same symbol-color combination indicates different subjects. The joint speeds of the same subject between small and large cylinders are connected by lines. Interspecies differences in DIP and MCP speeds cannot be accounted for by differences in the relative size of objects for large human hands vs. small monkey hands.

## References

Baldwin MKL, Cooke DF, Goldring AB, Krubitzer L (2018) Representations of fine digit movements in posterior and anterior parietal cortex revealed using long-train intracortical microstimulation in macaque monkeys. Cerebral Cortex 28:4244–4263.

Bernhard CG, Bohm E, Petersén I (1953) Investigations on the Organization of the Corticospinal System in Monkeys. Acta Physiologica Scandinavica 29:79–105 Available at: http://doi.wiley.com/10.1111/j.1365-201X.1953.tb10772.x.

Bird EE, Kivell TL, Skinner MM (2021) Cortical and trabecular bone structure of the hominoid capitate. Journal of Anatomy 239:351–373 Available at: https://onlinelibrary.wiley.com/doi/abs/10.1111/joa.13437 [Accessed July 30, 2021].

Brendel W, Romo R, MacHens CK (2011) Demixed principal component analysis. Advances in Neural Information Processing Systems 24: 25th Annual Conference on Neural Information Processing Systems 2011, NIPS 2011:1–9.

Castiello U, Dadda M (2019) A review and consideration on the kinematics of reach-to-grasp movements in macaque monkeys. Journal of Neurophysiology 121:188–204.

Christel MI, Billard A (2002) Comparison between macaques’ and humans’ kinematics of prehension: The role of morphological differences and control mechanisms. Behavioural Brain Research 131:169–184.

Diogo R, Wood BA (2012) Comparative Anatomy and Phylogeny of Primate Muscles and Human Evolution. Oxford: Taylor and Francis.

Durand J-B, Nelissen K, Joly O, Wardak C, Todd JT, Norman JF, Janssen P, Vanduffel W, Orban GA (2007) Anterior Regions of Monkey Parietal Cortex Process Visual 3D Shape. Neuron 55:493–505 Available at: http://www.sciencedirect.com/science/article/pii/S0896627307004990 [Accessed September 10, 2020].

Gharbawie OA, Stepniewska I, Qi H, Kaas JH (2011) Multiple parietal-frontal pathways mediate grasping in macaque monkeys. Journal of Neuroscience 31:11660–11677.

Goodman JM, Tabot GA, Lee AS, Suresh AK, Rajan AT, Hatsopoulos NG, Bensmaia S (2019) Postural Representations of the Hand in the Primate Sensorimotor Cortex. Neuron:566539 Available at: http://biorxiv.org/content/early/2019/03/05/566539.abstract.

Greenspon CM, Sobinov AR (2021) NCams. Available at: https://github.com/CMGreenspon/NCams [Accessed July 27, 2021].

Häger-Ross C, Schieber MH (2000) Quantifying the independence of human finger movements: comparisons of digits, hands, and movement frequencies. The Journal of neuroscience: the official journal of the Society for Neuroscience 20:8542–8550.

Heffner R, Masterton B (1975) Variation in form of the pyramidal tract and its relationship to digital dexterity. Brain, behavior and evolution 12:161–200 Available at: http://www.ncbi.nlm.nih.gov/pubmed/1212616.

Holzbaur KRS, Murray WM, Delp SL (2005) A model of the upper extremity for simulating musculoskeletal surgery and analyzing neuromuscular control. Annals of biomedical engineering 33:829–840 Available at: http://www.ncbi.nlm.nih.gov/pubmed/16078622.

Ingram JN, Körding KP, Howard IS, Wolpert DM (2008) The statistics of natural hand movements. Experimental Brain Research 188:223–236.

Iwaniuk AN, Pellis SM, Whishaw IQ (1999) Is digital dexterity really related to corticospinal projections?: A re-analysis of the Heffner and Masterton data set using modern comparative statistics. Behavioural Brain Research 101:173–187.

Jeannerod M (1988) The neural and behavioural organization of goal-directed movements. New York, NY, US: Clarendon Press/Oxford University Press.

Kaas JH, Qi H-X, Stepniewska I (2020) Evolution of Parietal-Frontal Networks in Primates. In: Evolutionary Neuroscience, pp 657–667. Elsevier. Available at: https://linkinghub.elsevier.com/retrieve/pii/B9780128205846000271.

Karakostis FA, Haeufle D, Anastopoulou I, Moraitis K, Hotz G, Tourloukis V, Harvati K (2021) Biomechanics of the human thumb and the evolution of dexterity. Current Biology 0 Available at: https://www.cell.com/current-biology/abstract/S0960-9822(20)31893-5 [Accessed March 5, 2021].

Kivell TL (2016) The Primate Wrist. In: The Evolution of the Primate Hand (Kivell TL, Lemelin P, Richmond BG, Schmitt D, eds). New York: Springer Science.

Kivell TL, Kibii JM, Churchill SE, Schmid P, Berger LR (2011) Australopithecus sediba Hand Demonstrates Mosaic Evolution of Locomotor and Manipulative Abilities. Science 333:1411–1417 Available at: https://science.sciencemag.org/content/333/6048/1411 [Accessed July 30, 2021].

Kivell TL, Lemelin P, Richmond BG, Schmitt D eds. (2016) The Evolution of the Primate Hand. New York, NY: Springer New York. Available at: http://link.springer.com/10.1007/978-1-4939-3646-5.

Kobak D, Brendel W, Constantinidis C, Feierstein CE, Kepecs A, Mainen ZF, Qi XL, Romo R, Uchida N, Machens CK (2016) Demixed principal component analysis of neural population data. eLife 5:1–36.

Lemon R (2019) Recent advances in our understanding of the primate corticospinal system. F1000Research 8 Available at: http://www.ncbi.nlm.nih.gov/pubmed/30906528.

Marzke MW (2013) Tool making, hand morphology and fossil hominins. Philosophical Transactions of the Royal Society B: Biological Sciences 368:20120414 Available at: https://royalsocietypublishing.org/doi/full/10.1098/rstb.2012.0414 [Accessed July 30, 2021].

Mathis A, Mamidanna P, Cury KM, Abe T, Murthy VN, Mathis MW, Bethge M (2018) DeepLabCut: markerless pose estimation of user-defined body parts with deep learning. Nat Neurosci 21:1281–1289.

Mayer A, Baldwin MKL, Cooke DF, Lima BR, Padberg J, Lewenfus G, Franca JG, Krubitzer L (2019) The Multiple Representations of Complex Digit Movements in Primary Motor Cortex Form the Building Blocks for Complex Grip Types in Capuchin Monkeys. The Journal of neuroscience: the official journal of the Society for Neuroscience 39:6684–6695.

Mountcastle VB, Lynch JC, Georgopoulos A, Sakata H, Acuna C (1975) Posterior parietal association cortex of the monkey: command functions for operations within extrapersonal space. Journal of Neurophysiology 38:871–908 Available at: https://journals.physiology.org/doi/abs/10.1152/jn.1975.38.4.871 [Accessed April 4, 2021].

Napier JR (1967) Hands (Tuttle RH, ed). Princeton, New Jersey: Princeton University Press.

O’Connor DH, Krubitzer L, Bensmaia S (2021) Of mice and monkeys: Somatosensory processing in two prominent animal models. Progress in Neurobiology:102008.

Okorokova E V., Goodman JM, Hatsopoulos NG, Bensmaia SJ (2019) Decoding hand kinematics from population responses in sensorimotor cortex during grasping.

Patel BA, Maiolino SA (2016) Morphological Diversity in the Digital Rays of Primate Hands. In: The Evolution of the Primate Hand.

Penfield W, Rasmussen T (1950) The cerebral cortex of man: a clinical study of localization of function. New York: Macmillan.

Rathelot JA, Strick PL (2009) Subdivisions of primary motor cortex based on cortico-motoneuronal cells. Proceedings of the National Academy of Sciences of the United States of America 106:918–923.

Rolian C, Lieberman DE, Zermeno JP (2011) Hand biomechanics during simulated stone tool use. Journal of Human Evolution 61:26–41 Available at: https://www.sciencedirect.com/science/article/pii/S0047248411000492 [Accessed July 30, 2021].

Sakata H, Taira M, Murata A, Mine S (1995) Neural Mechanisms of Visual Guidance of Hand Action in the Parietal Cortex of the Monkey. Cereb Cortex 5:429–438 Available at: https://academic-oup-com.proxy.uchicago.edu/cercor/article/5/5/429/375668 [Accessed September 1, 2020].

Santello M, Flanders M, Soechting JF (1998) Postural Hand Synergies for Tool Use. J Neurosci 18:10105–10115.

Santello M, Soechting JF (1998) Gradual Molding of the Hand to Object Contours. Journal of Neurophysiology 79:1307–1320.

Saul KR, Hu X, Goehler CM, Vidt ME, Daly M, Velisar A, Murray WM (2015) Benchmarking of dynamic simulation predictions in two software platforms using an upper limb musculoskeletal model. Computer methods in biomechanics and biomedical engineering 18:1445–1458 Available at: http://www.ncbi.nlm.nih.gov/pubmed/24995410.

Schaffelhofer S, Agudelo-Toro A, Scherberger H (2015) Decoding a wide range of hand configurations from macaque motor, premotor, and parietal cortices. Journal of Neuroscience 35:1068–1081.

Schaffelhofer S, Scherberger H (2016) Object vision to hand action in macaque parietal, premotor, and motor cortices Kastner S, ed. eLife 5:e15278 Available at: https://doi.org/10.7554/eLife.15278 [Accessed September 9, 2020].

Schieber MH (1991) Individuated finger movements of rhesus monkeys: A means of quantifying the independence of the digits. Journal of Neurophysiology 65:1381–1391.

Schieber MH, Santello M (2004) Hand function: peripheral and central constraints on performance. Journal of Applied Physiology 96:2293–2300.

Schilling A-M, Tofanelli S, Hublin J-J, Kivell TL (2014) Trabecular bone structure in the primate wrist. Journal of Morphology 275:572–585 Available at: https://onlinelibrary.wiley.com/doi/abs/10.1002/jmor.20238 [Accessed July 30, 2021].

Seth A, Hicks JL, Uchida TK, Habib A, Dembia CL, Dunne JJ, Ong CF, DeMers MS, Rajagopal A, Millard M, Hamner SR, Arnold EM, Yong JR, Lakshmikanth SK, Sherman MA, Ku JP, Delp SL (2018) OpenSim: Simulating musculoskeletal dynamics and neuromuscular control to study human and animal movement Schneidman D, ed. PLoS Comput Biol 14:e1006223.

Sobinov AR, Bensmaia SJ (2021) The neural basis of manual dexterity. Nature Reviews Neuroscience.

Stepniewska I, Gharbawie OA, Burish MJ, Kaas JH (2014) Effects of muscimol inactivations of functional domains in motor, premotor, and posterior parietal cortex on complex movements evoked by electrical stimulation. Journal of Neurophysiology 111:1100–1119.

Strick PL, Dum RP, Rathelot J-A (2021) The Cortical Motor Areas and the Emergence of Motor Skills: A Neuroanatomical Perspective. Annual Review of Neuroscience 44:null Available at: https://doi.org/10.1146/annurev-neuro-070918-050216 [Accessed April 22, 2021].

Taira M, Mine S, Georgopoulos AP, Murata A, Sakata H (1990) Parietal cortex neurons of the monkey related to the visual guidance of hand movement. Exp Brain Res 83:29–36.

Thakur PH, Bastian AJ, Hsiao SS (2008) Multidigit movement synergies of the human hand in an unconstrained haptic exploration task. Journal of Neuroscience 28:1271–1281.

Todorov E, Ghahramani Z (2004) Analysis of the synergies underlying complex hand manipulation. In: The 26th Annual International Conference of the IEEE Engineering in Medicine and Biology Society, pp 4637–4640.

Tunik E, Frey SH, Grafton ST (2005) Virtual lesions of the anterior intraparietal area disrupt goal-dependent on-line adjustments of grasp. Nature Neuroscience 8:505–511.

Williams EM, Gordon AD, Richmond BG (2010) Upper limb kinematics and the role of the wrist during stone tool production. American Journal of Physical Anthropology 143:134–145 Available at: https://onlinelibrary.wiley.com/doi/abs/10.1002/ajpa.21302 [Accessed July 30, 2021].

Williams-Hatala EM, Hatala KG, Key A, Dunmore CJ, Kasper M, Gordon M, Kivell TL (2021) Kinetics of stone tool production among novice and expert tool makers. American Journal of Physical Anthropology 174:714–727 Available at: https://onlinelibrary.wiley.com/doi/abs/10.1002/ajpa.24159 [Accessed July 29, 2021].

Yan Y, Goodman JM, Moore DD, Solla SA, Bensmaia SJ (2020) Unexpected complexity of everyday manual behaviors. Nature Communications 11:1–8.

Young RW (2003) Evolution of the human hand: The role of throwing and clubbing. Journal of Anatomy 202:165–174.

